# Benchmarking Polygenic Risk Score Model Assumptions: towards more accurate risk assessment

**DOI:** 10.1101/2022.02.18.480983

**Authors:** Scott Kulm, Jason Mezey, Olivier Elemento

## Abstract

Polygenic risk scores represent an individual’s genetic susceptibility to a phenotype. Like with any models, statistical models commonly employed to fit polygenic risk scores and assess their accuracy contain several assumptions. The effects of these assumptions on models of polygenic risk score have not been thoroughly assessed.

We assessed 26 variations of the traditional polygenic risk score model, each of which mitigate assumptions in one of five facets of disease modelling: representation of age (6 variations), censorship (3 variations), competing risks (7 variations), formation of disease labels (6 variations), and selection of covariates (4 variations). With data from the UK Biobank, each model variation included age, sex, and a polygenic risk score derived from the PGS Catalog. Each of the 26 model variations were fitted to predict 18 diseases. Compared to the plain model that contained all five facets of assumptions, the model variations often fit the data better and generated predictions that largely differed from the predictions of the plain model. The statistic Royston’s R^2^ measured a model’s goodness of fit, and thereby determined if the model was an enhancement upon the plain model. For 15 of the 26 model variations Royston’s R^2^ was greater than that of the plain model for >50% of diseases. Reclassification rates, defined as the fraction of individuals in the top five percentiles of the plain model’s predictions who are not in the top five percentiles of a model variation’s predictions, was used to determine if the variation led to significantly different predictions. For 20 of the 26 model variations the median reclassification rate calculated across the 18 diseases was greater than 10%. Comparisons of accuracy statistics further illustrated how much each model variation’s predictions differed from the plain model’s predictions.

Models containing polygenic risk scores appear to be significantly affected by many common modelling assumptions. Therefore, future investigations should consider taking some action to mitigate modelling assumptions.

**Author Summary:** An individual’s genetics can increase their risk of experiencing a disease. The exact magnitude of the increased risk is estimated within a statistical model. The traditional model type employed in this process is relatively plain and contains several assumptions. The predicted risk estimates from this plain model may be unnecessarily inaccurate. To test this possibility, we searched the literature for model variations that reduce the assumptions of the plain model, ultimately creating 26 distinct model variations that may improve upon the plain model. Each model variation was fit with data from the UK Biobank to predict 18 diseases. We found that 15 of the 26 models variations fit the data better than the plain model for a majority of diseases. Goodness of fit was measured with Royston’s R^2^ statistic. Further calculations found that the predictions of the model variations were often significantly more or less accurate than the predictions of the plain model. We believe these results indicate that future investigations of polygenic risk scores should not employ the plain model, as unreliable risk predictions will likely result.

## Introduction

There is growing interest in leveraging polygenic risk scores as a way of estimating an individual’s genetic risk of developing certain diseases. Initially introduced in 2009 [1], and quickly grown in popularity [1, 2], polygenic risk scores are defined as the sum of risk alleles which are weighted by effect sizes determined from genome wide association studies [3]. The resulting score can, in theory, be combined with other clinical metrics and risk factors such as age or body mass index (BMI) to form a complete disease risk prediction framework [4, 5].

Within a precision medicine paradigm, such predictions may eventually spur beneficial clinical actions, e.g. frequent disease monitoring, early detection or prophylactic measures, and thereby improve patient outcomes [6, 7].

The contribution of a polygenic risk score to an estimation of disease risk is determined through modelling. Most models employed to estimate the importance of polygenic risk scores logistically regress the polygenic risk score alongside a few covariates, such as age and sex, against disease labels usually obtained via electronic health records [8, 9]. While this model is inherently wrong [10], it does provide some use as it combines a polygenic risk score with other risk factors into a single risk prediction. Some models employed to estimate a polygenic risk score’s accuracy are termed survival models, as their dependent variable is the duration of time until a disease of interest occurs. These models generally censor individuals after exiting the study, and, as we further discuss below, often do not consider health events that may compete with the primary disease of interest and thus change outcomes. For example, the outcome of death competes with, and thereby hinders, the observation of an Alzheimer’s disease diagnosis.

There are a range of approaches previously published in other fields that can be used to mitigate this problem, which we explore here. Additional assumptions are made within a typical polygenic risk score survival model. How all of these assumptions impact a model’s goodness of fit and generated predictions have not been tested, to the best of our knowledge. We therefore consider five specific areas in which a traditional, plain polygenic risk score model contains assumptions or biases. These five areas are as follows.

First, in the vast majority of polygenic score models, age, which frequently influences disease risk [11–13], is only considered through the inclusion of a single covariate in a linear model. While the implied assumption that age has a linear relationship with disease risk is dubious in general [11], it is especially improbable in relation to polygenic risk scores, which are more accurate at predicting early onset, as compared to late onset, disease [14, 15]. To better model the complexities of age two categories of variations are available. First, nonlinear age covariates, such as age squared, can be added to the model [16, 17] Second, the cohort can be altered, for example the age distribution of individuals without disease can be made to match the age distribution of individuals with disease [18]. While both categories of modeling variations are made in GWAS studies [16, 17], they are rarely considered, to the best of our knowledge, within polygenic risk score analyses.

Second, the time in which each individual survives without disease is quantified by the number of days since assessment. This approach creates an immortality bias within the individuals who were assessed at the end of the enrollment period, since they were unable to experience the disease during a period of time in which most other individuals in the study could[19]. The bias can be reduced by recalculating each individual’s survival time in reference to a fixed date, forming a delayed entry model [20]. Alternatively, each individual’s survival time can be calculated in reference to their date of birth [21, 22]. While models with delayed entry have been implemented in the wider epidemiological field, they have not to our knowledge been employed in conjunction with polygenic risk scores.

Third, standard cox proportional hazard models assume that all individuals will eventually be diagnosed with the disease, a poor approximation considering many individuals will die before the disease even begins to develop [23]. The implementation of well-established competing risk models, such as multi-state or Fine and Gray, provide a ready correction to this assumption [24]. Moreover, correcting for competing risks leads to absolute risk estimates, which are preferred within current real-world polygenic risk score implementations as they are easier to interpret in a clinical context [25]. Despite the preference towards absolute risk estimates, the vast majority of models do not consider competing risks.

Fourth, the disease labels that critically determine the accuracy of risk predictions are commonly based on unreliable sources. Electronic health records can be miscoded or partially unavailable due to a change in primary health provider [26], and self-reported records can be liable to an individual’s memory or perceptions [27]. The problem of unreliable disease labels is largely irremediable using current data collection mechanisms. However, patient data, such as HbA1c levels, blood pressure and BMI, can be used to predict a disease’s risk, such as Type II Diabetes.

If a prediction derived from the patient data confidently disagrees with the disease label derived from electronic health records, then the corresponding suspect individual could be removed from the analysis. Previous studies have demonstrated with evidence the accuracy of disease predictions derived from large datasets of patient data [28, 29], but to the best of our knowledge they have not assessed whether using disease predictions instead of direct disease labels effects the predictions of polygenic risk score models.

Fifth, the features/covariates included in disease prediction models alongside the polygenic risk score are often manually and sparingly chosen. In this situation the polygenic risk score may become confounded, and thereby improperly represent an individual’s genetic risk. For example, a model of heart disease that includes the covariates of age, sex, BMI and family history may seem to contain all relevant features, yet blood pressure and/or smoking history may also be relevant to the prediction of heart disease [30, 31]. The application of a systematic feature selection process could unbiasedly determine which features should be included in a model so that the polygenic risk score is properly adjusted. While such processes do exist, to our knowledge nearly all investigations manually select a limited set of model covariates.

These five areas in which polygenic risk score models make assumptions and simplifications all correspond to known mitigating model variations. An objective way of determining whether the variations are empirically better models is goodness of fit statistics. If a model variation fits the data better than the plain model, then we considered it to be an enhancement to the plain model which should likely be preferentially employed. Although polygenic risk score models are ultimately intended to predict the relative risk of an individual with respect to a given disease. Goodness of fit statistics are unable to validate the accuracy of these predictions. Therefore, along with determining whether a model variation is relatively enhanced, we also calculated accuracy statistics to better understand exactly how each assumption affects the model’s predictions. We decided to report the accuracy statistics of both enhanced and non-enhanced model variations, because the non-enhanced models could still possibly correct for an improper bias in the data [32–35]. For example, a model that is massively confounded may be more accurate than a model that properly controls for confounders, but that does not imply that the confounded model should be preferred or considered more useful. We determined the models that are likely to reduce such biases, and thereby deserve reporting, by their level of support in the literature. While these model variations are common in certain areas of epidemiology, to our knowledge they have seen little use within the vast majority of polygenic risk score analyses.

Little is therefore known concerning the practical effect of each variation. Following claims made in the literature, we hypothesized that variations to the plain, traditional model containing all five facets of assumptions often led to enhanced models whose predictions significantly differed from the predictions of the plain model. If these predictions were eventually implemented in the clinic, significant differences in predictions would lead to a significant proportion of individuals experiencing different clinical actions. If these revised clinical actions were informed by the predictions of the enhanced mode, then on average, we could expect better outcomes.

We tested our hypothesis by first acquiring data from UK Biobank and PGS Catalog (Figure 1A). Second, we processed the data, forming polygenic risk scores, predictions from CoxNet models which were fit on a dataset containing many diverse features, and phenotypes derived from electronic health records (Figure 1B). Third, we addressed one of five areas in which the plain, traditional model makes assumptions or holds biases by forming 26 variations of the plain model (Figure 1C). Each model variation was fit to 18 diseases. The covariates of each model included age, sex, top ten genetic principal components and a disease-specific polygenic risk score derived from the PGS Catalog. Each model’s goodness of fit was measured with pseudo R^2^ values. If the R^2^ value of a model variation was greater than the R^2^ value of the plain model for a majority of diseases, then we considered the model variation an enhanced method of fitting the data. The models predictions, made upon a withheld testing dataset, formed accuracy statistics, such as concordance and odds ratio. These accuracy statistics were compared in two ways (Figure 1D). First, we compared the statistics from a model variation to the statistics from a plain model that approximates the approach taken by most polygenic risk score analyses. The resulting difference in these statistics was termed the statistical difference. Second, we compared the statistics from a model with a polygenic risk score to the statistics from the same model but with the polygenic risk score removed. The resulting difference in these statistics was termed the statistical improvement, which highlights how the model’s predictions change when a polygenic risk score is additionally considered. Difference and improvement statistics were combined to determine the impact of the model variation on the relative contribution of the polygenic risk score upon the model’s predictions. Goodness of fit metrics generated from the model variations are not necessarily greater than the goodness of fit metrics generated from the plain model. However, we still considered and reported on all the model variations because the literature strongly supports each model variation’s ability to eliminate assumptions or biases.

**Figure 1:**
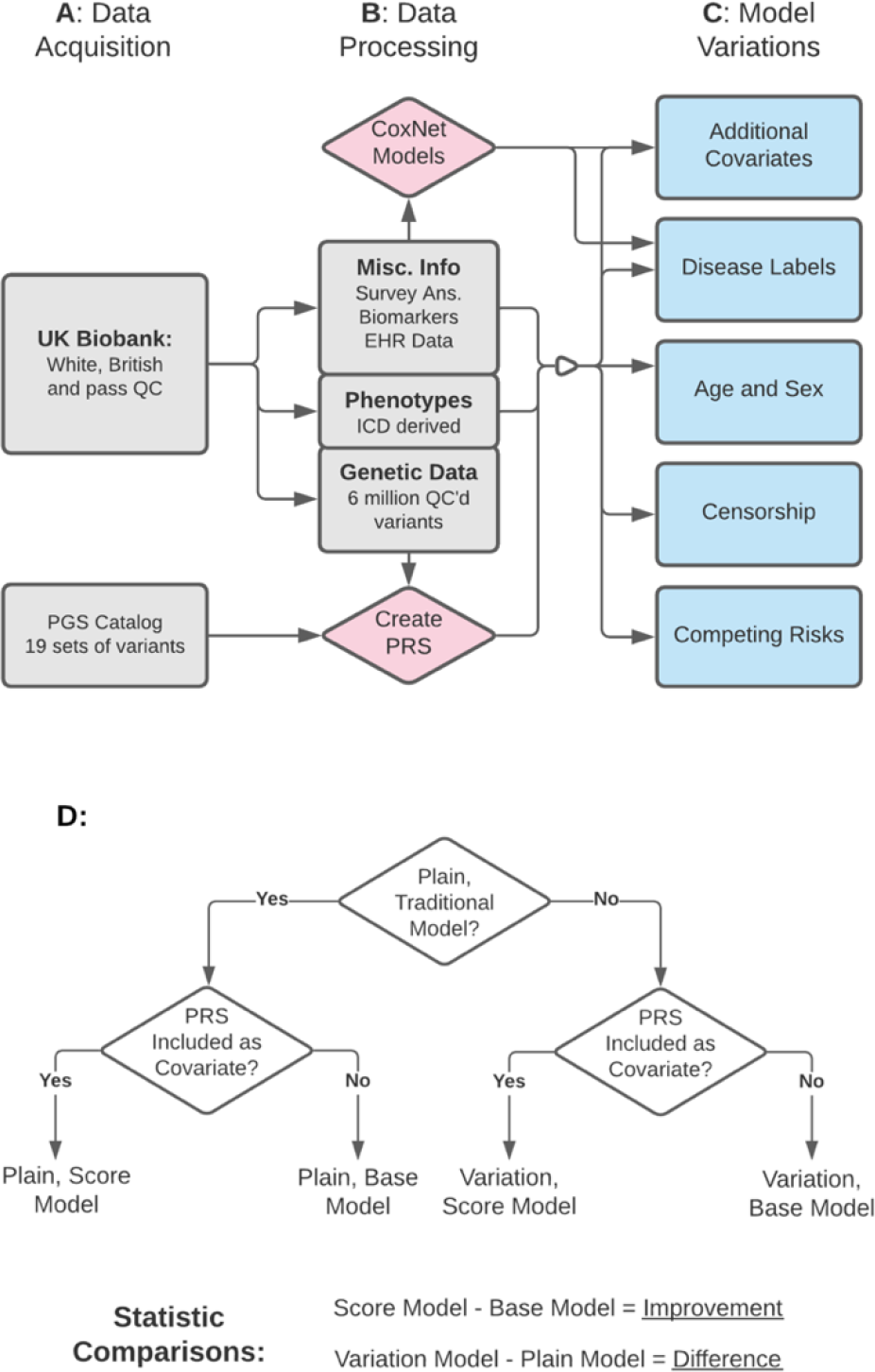
Methodological Flow Chart. A: All possible data from the UK Biobank was reduced to a homogenous, quality-controlled population then split into like data types. Sets of variants were acquired from the PGS Catalog. B: Genetic data created polygenic risk scores and miscellaneous data formed non-genetic disease predictions within CoxNet models. C: For each of five distinct areas of modelling assumptions, model variations formed predictions upon the testing data which were converted into accuracy statistics. D: Specific terms are employed to describe models and statistics. A plain model utilizes the plain framework which is a cox proportional hazard model that regresses the covariates of age, sex and top ten genetic principal components against right-censored survival times. Model variations make some modification to the plain model with the goal of removing assumptions or biases. Score models contain a polygenic risk score whereas as base models do not. Improvement statistics subtract a statistic generated from a score model from a statistic generated from a base model. Difference statistics subtract a statistic from an enhanced model from a statistic generated from a plain model. Both difference and improvement statistics are only computed between models of the same disease.

The results from these comparisons supported our hypothesis that many of the variations made to the plain model led to enhanced models whose predictions differed significantly from the plain model’s predictions.

## Results

### Characterization of Plain Disease Prediction Models Which Contain Polygenic Risk Scores

To test our hypothesis that attempts to reduce the assumption made within the plain model leads to the formation of enhanced models whose predictions vary greatly from the predictions of the plain model, we first characterized the plain model when applied to each of the 18 diseases of interest. This plain model type was specifically a cox proportional hazard model that regressed a small set of covariates against a right-censored time to disease. A plain model type which included the covariate of age, sex, top ten genetic principal components and a polygenic risk score was specifically termed a plain, score model. The polygenic risk score term in every disease’s plain, score model was significant after Bonferroni correction (median P = 3.02e-39) (Table S1.1). The goodness of fit of the plain model was measured by Royston’s pseudo-R^2^, which is calculated from the commonly used Royston and Sauerbrei’s D statistic [36–38]. The average R^2^ was 0.151 (SE 0.0221). The minimum R^2^ was 0.0166 for breast cancer and the maximum R^2^ was 0.333 for gout (Table S1.1). These values will be later compared to the R^2^ values for the variation models to determine if they serve as model enhancements.

**Table 1:** Summary of All Variation Models. The model variation column describes models from each of the five areas of modelling assumptions. The description column indicates how exactly the model variation differs from the plain model. The column of average R^2^ difference was determined by first calculating Royston’s pseudo R^2^ for all model variations, then subtracting the R^2^ of the model variation from the R^2^ of the plain model for each disease. This R^2^ difference is averaged across diseases, resulting in the value indicated. The column diseases with positive R^2^ difference was determined by counting the diseases in the preceding calculation whose R^2^ difference was positive.

| Model Variation | Description | Average R<sup>2</sup> Difference | Diseases with Positive R<sup>2</sup> Difference |
| --- | --- | --- | --- |
| Age |  |  |  |
| Squared | age<sup>2</sup> added to model | 0.00746 | 18 |
| Interaction | age<sup>2</sup> and age:PRS added to model | 0.00897 | 18 |
| Date of Birth | Current age replaces age at analysis | -0.006 | 2 |
| Age Stratified | Stratified analysis by 5 year age groups | -0.0433 | 5 |
| Narrow Age | Only individuals with age 56-64 | -0.0620 | 3 |
| Age Match | Age distribution of cases matched by controls | -0.092 | 0 |
| Censor |  |  |  |
| No censoring | All individuals end follow-up at present | -0.0188 | 6 |
| Delay Entry by Study Time | Counting days in study, starting at 1/1/2006 | 0.00147 | 12 |
| Delay Entry by Age Time | Counting days alive, starting at date of birth | -0.0921 | 0 |
| Competing Risk |  |  |  |
| Fine-Gray | Fine-Gray model employed with death competing against disease | -6.08E-6 | 11 |
| Multi-State | Multi-State model employed with death competing against disease | -0.0149 | 6 |
| Risk Regression | Risk Regression model employed with death competing against disease | -1.974E-4 | 10 |
| iCARE: Holt | Absolute risk model fit with iCARE package; disease incidence estimated by UKBB with Holt smoothing | -8.45E-12 | 1 |
| Disease Labels |  |  |  |
| Reclassify 5% | Individuals in top/bottom 5% according to CoxNet predictions with negative/positive label are reclassified | 0.0973 | 17 |
| Remove 5% | Individuals in top/bottom 5% according to CoxNet predictions with negative/positive label are removed | 0.0259 | 18 |
| Additional Covariates |  |  |  |
| Incl. CoxNet Pred. | CoxNet Pred. Added to model | 0.312 | 18 |
| Incl. Top 5 Feats. | Five features with greatest absolute coefficient in CoxNet model added to model | 0.250 | 18 |

Predictions from plain, score models were used to compute concordances, which are a measure of model accuracy that consider all individuals. A concordance of 0.0 indicates a perfectly inaccurate prediction and a concordance of 1.0 represents a perfectly accurate prediction. The average concordance of the plain, score models was 0.688 (SE 0.0126). To estimate the value a polygenic risk score provides when it is added to a disease prediction model we constructed plain, base models, which also employed the simple model type but only included the covariates of age, sex, and top ten genetic principal components. The concordance of a plain, score model minus the concordance of a plain, base model was termed concordance improvement. The average concordances improvement was 0.044 (SE 0.0130) (Table S1.3). The difference between an acceptable and good concordance, and the difference between a good and excellent concordance is often assumed to be 0.1 [39, 40]. Therefore, on average, the addition of a polygenic risk score to a base model makes a fair predictor approximately halfway closer to becoming a good predictor.

To determine if the accuracy indicated by the concordance statistic is shared by other accuracy metrics, we repeated many of the comparisons, with the same terminology, but instead of concordance calculated odds ratios. We computed odds ratios from a 2x2 contingency table in which the high-risk, or exposed group, are the individuals in the top 5 percentiles of a model’s predictions and the low-risk, or unexposed group, are the individuals in the bottom 50 percentiles of a model’s predictions. The log odds ratios generated from plain, score models were on average 1.82 (SE 0.138). The log odds ratio improvement, defined as the difference between the log odds ratio generated by a plain, score model and the log odds ratio generated by a plain, base model, averaged 0.482 (SE 0.145) (Table S1.5). We report odds ratios on the log scale because doing so provides numerous beneficial properties: symmetric error bars, an underlying normal distribution, and a range of possible values than span from negative to positive infinity [41]. The log odds ratio can by exponentiated at anytime to generate more interpretable values. For example, a log odds ratio of 1.82 is an odds ratio of 6.17, which implies that the odds of having the disease in the high-risk group is 517% larger than the odds of having the disease the disease in the low-risk group. The same types of statistics and terminology used to assess the plain model were also applied to each of the five enhanced analyses. In addition, we computed difference statistics and reclassification rates. Differences statistics were defined as a statistic generated from a variation model minus a statistic generated from a plain model.

Reclassification rates are calculated as the fraction of individuals who were considered as high-risk according to a plain model’s predictions but are no longer classified as high-risk according to a variation model’s predictions. These difference statistics and reclassification rates provided a direct means of testing our hypothesis.

### Modified Covariate Representation of Age improves PRS accuracy and goodness-of-fit

We hypothesized that models with modified representation of age would better fit the data and generate predictions that substantially differ from the predictions of the plain models. The plain model represents age as a single covariate. Model variations that modify the representation of age either included additional covariates or changed the cohort. For example, the age squared model added the covariate of age squared, whereas the narrow age cohort model limited the cohort to individuals between the age of 55 and 64. A list of other variations made to the plain model alongside their descriptions are provided in Table 1, with additional information located in the supplemental methods.

We first evaluated whether the model variations that modified the representation of age were enhancements to the plain model by comparing Royston’s pseudo-R^2^ values. The two model variations that added covariates to the plain model, squared and interaction, were both definitive enhancements. Specifically, the squared model variation added the covariate age squared to the plain model and the interaction model variation added the covariates age squared and an interaction between age and the polygenic risk score to the plain model. The R^2^ of both the squared and interaction model was greater than the R^2^ of the plain model for all diseases (Table 1). We term the difference between a model variation’s R^2^ and a plain model’s R^2^ the R^2^ difference. If the R^2^ differences were positive for over 50% of the diseases under a model variation, then we consider the model variation to be an enhancement upon the plain model.

None of the model variations that altered the cohort (age stratified, narrow age, age match) were enhancements upon the plain model. The average R^2^ that corresponded to each of these model variations was negative (Table 1, S2.1). The decreased sample size of these models may partially explain why they cannot be considered enhancements. Lastly, the date of birth model variation, which replaced the plain model’s age covariate measured at time of the analysis with age at birth, was also unlikely to enhance the plain model as its average R^2^ was negative (Table 1, S2.1).

Goodness of fit statistics thereby identify two of the sex model variations that modified representation of age as enhanced models.

We next compared the predictions of each model that modified age representation to the predictions of the plain model through the calculation of reclassification rates. Reclassification rates are defined as the proportion of a high-risk group that changes when one set of model predictions is replaced by another set of model predictions. The model predictions used to define the high-risk group could be extracted at any time-point in the study (Figure 2A). Fortunately, no matter the time-point chosen, the individuals in the high-risk group will be identical, greatly simplifying the statistic calculations. The resulting reclassification rates for each age modified model varied greatly across diseases. The age stratified model, which modeled sub-cohorts that contained individuals of similar ages then combined the results, generated the highest average reclassification rate at 43.4% (SE 5.1%), with values for each disease ranging widely from only 10.2% for schizophrenia to 71.4% for gout. The squared model, which adds an age squared term to the plain model, generated the lowest average reclassification rate of 6.4% (SE 1.4%), with a maximum reclassification rate of 20.7% for prostate cancer and a minimum of 0.1% for celiac disease (Figure 2B, Table S2.4). While reclassification rates directly exhibit how the differences between predictions from the model variation and plain model vary greatly across diseases, they are unable to express the accuracy of the predictions.

**Figure 2:**
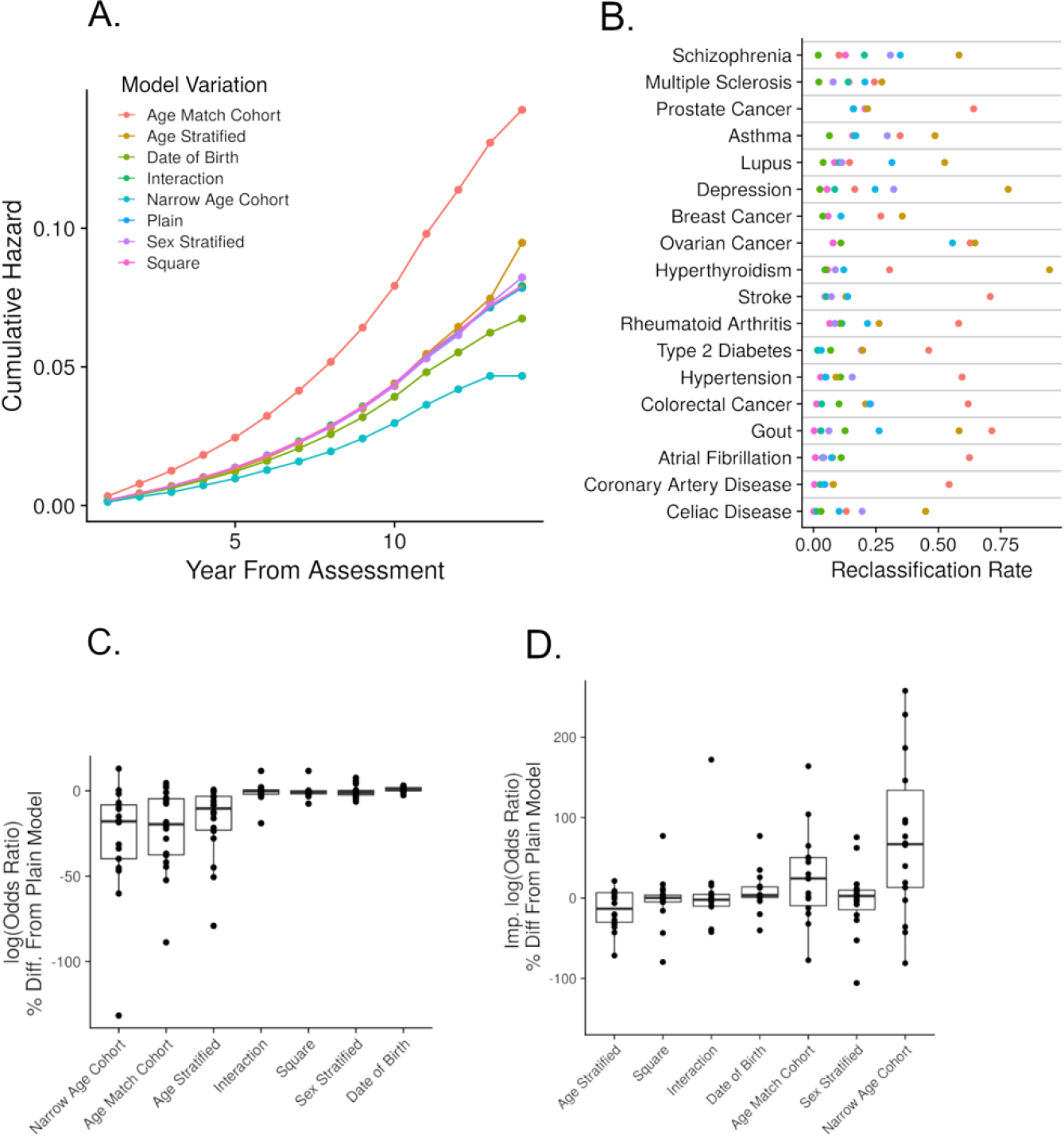
Model Variations that Modify the Representation of Age. A: Kaplan-Meier style curve of cumulative hazard predictions for stroke, estimated by various cox proportional hazard models. B: For each disease, the reclassification rates, or fractions of individuals who are no longer considered to be in the top 5% of risk when replacing the plain model prediction with a model variation prediction. C: For each model the percent difference between a model variation and plain model’s odds ratio. Odds ratios are calculated by comparing those in the bottom half of the model’s predictions against those in the top 5%. D: Same comparison as in C, except the odds ratio improvement, or the difference in odds ratio computed from the score and base model, replaced the odds ratio directly computed from the score model.

Odds ratios provide a simple method of determining how accurate model predictions are. Odds ratios were specifically computed from a 2x2 contingency table in which the high-risk, or exposed, group was in the top 5% of a score model’s predictions and the low-risk, or unexposed, group was in the bottom 50%. The log odds ratio of a model variation minus the log odds ratio of the plain model generated a difference log odds ratio. The three model variations that generated the greatest average log odds ratio differences from the plain model were the narrow-age (-27.9%, SE 7.6%), age-matched (-22.6%, SE 5.8%) and age-stratified (-17.7%, SE 5.3%). The age-matched model alters the cohort such that the distribution of age in the cases matches the distribution of age in the controls, and the narrow-age model only considers individuals aged 56-64. All three of these analyses directly alter the composition of the cohort under model.

Comparatively, the average log odds ratio differences of the squared model (0.66%, SE 0.85%) and interaction model (1.0%, SE 1.4%) were relatively small (Fig. 2C, Table S2.5). Both of these models modify age’s representation through the addition of model covariates.

Odds ratios reflect the effect of all involved covariates, not just the polygenic risk score. To get a better sense of how well the polygenic risk score alters the model predictions, for a single disease we subtracted the score model’s log odds ratio from the base model’s log odds ratio. We termed this comparison log odds ratio improvement. The two analyses with the greatest average improved log odds ratio are narrow age (0.62, SE 0.13) and age-match (0.57, SE 0.14), both of which change the cohort under analysis. The two analyses with the lowest averaged improved log odds ratios are square (0.50, SE 0.14) and interaction (0.53, SE 0.14), both of which add covariates to the plain model. The improved log odds ratio of the narrow age model was on average 104% (SE 42%) greater than the improved log odds ratio of the plain model, suggesting that the contribution of a polygenic risk score to the disease model may be more than double what is commonly estimated in a plain model (Fig. 2D, Table S2.5). Models that modify the representation of age through alterations of the cohort under analysis generate predictions that are less accurate than the plain model but are more effected by the polygenic risk score.

### Modifications to the Manner Individuals are Censored improve PRS goodness-of-fit

We hypothesized that modifications to the way individuals are censored within the plain model would lead to enhanced model variations whose predictions significantly differ from the plain model’s predictions. The plain model tracks the time of each individual in the study, starting at the date of assessment. This approach gives rise to an immortality bias, as individuals that enter the study at a relatively later dates have shorter time frames in which they may possibly experience the disease of interest [19]. Even greater biases can occur when censorship does not occur, an approximation of the popular logistic regression. These biases can potentially be diminished by delaying the time of entry into the study. Individuals can enter the study at a time value measured in reference to either a fixed date, forming the delay entry at study time model, or their date of birth, forming the delay entry at age time model (Figure 3A, Table 1, M1).

**Figure 3:**
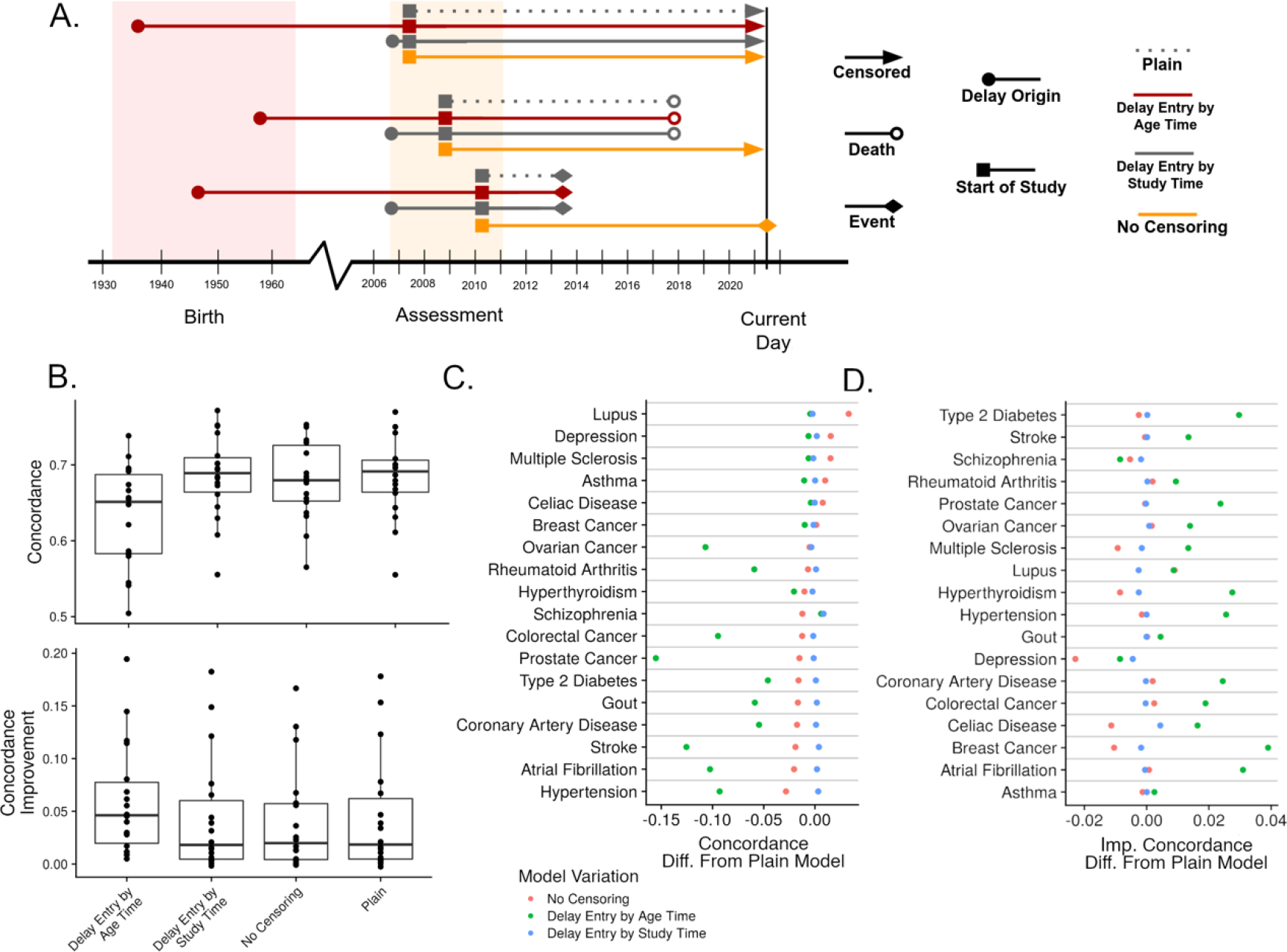
Model Variations that Modify the Manner Individuals are Censored. A: Diagram of three hypothetical individuals within the cohort under analysis. The different color and line types indicate the different model variations. B: The concordance and concordance improvement stratified by model variation. Concordance improvement is the difference in concordances calculated from score and base models. C: For each disease the score model’s concordance computed from a model variation minus the concordance computed from a plain, score model. D: Same comparison as in C but concordance improvement, or the difference in score and base model concordances, is displayed.

Variations to the manner the plain model censor individuals led to models that were enhanced according to goodness of fit measurements. Specifically, the difference between the delay entry by study time model’s R^2^ and the plain model’s R^2^ was positive for 12 diseases (Table 1, S3.1). The two other variations to the plain model that involved censorship did not lead to enhanced models. The delay entry by age time model’s R^2^ values were less than plain model’s R^2^ for all diseases. The no censorship model, which was expected to perform poorly, interestingly generated R^2^ values greater than those of the plain model for 6 diseases (Table 1, S3.1). These 6 diseases, which include asthma, depression and schizophrenia, are not particularly known for occurring near the end of life [42–44], which may explain why they are fit well by the no censorship model. We therefore found 1 model variation that enhanced the plain model by modifying the manner in which individuals were censored.

Predictions generated from models with modified approaches to censoring were not substantially different than predictions generated from plain models. The concordances of the no censorship models were on average 0.73% (SE 0.54%) less than the concordances of the plain models. The concordances of the delay entry by study-time model were on average 0.08% (SE 0.09%) less than the concordances of the plain models. The concordances of a third approach which also employed delayed entry but shifted age from an explicit covariate to the timescale, were on average 7.60% (SE 1.70%) less than the concordances of the plain model (Figure 3B). The diseases with the largest concordance differences between the plain and delayed entry by age include prostate cancer (-22.3%) and stroke (-17.7%), both of which are particularly affected by age [45, 46] (Figure 3C, Table S3.3). Other diseases that are minimally affected by age did not generate a large concordance difference, suggesting that censorship approaches may alter prediction accuracies by modifying the effect of age.

Closer examination of concordance values indicated that modifications to the manner individuals are censored increases the amount the polygenic risk score contributed to the model. The average difference in concordance improvement from the no censoring model to the plain model is 1.96% (SE 7.21%). Similarly, the average difference in concordance improvement from the delay entry by study time model to the plain model to the is 3.00% (SE 2.02%). However, a much larger average difference in concordance improvement was computed between the delay entry by age time model and plain model, 143% (SE 67.7%) (Figure 3B). The difference in concordance improvement between the delay entry by age time model and the plain model is particularly large for gout (840%) and rheumatoid arthritis (429%) – both diseases that experience large age effects. Although, these differences reflect only modest absolute concordance improvements for the delay entry by age time model, 0.0050 for gout and 0.0116 for rheumatoid arthritis [47, 48] (Figure 3D, Table S3.3). Therefore, censorship approaches likely acting through modifications to the effect of age, have limited yet definite potential to alter the contribution of a polygenic risk score upon a disease model’s predictions.

### Consideration of Competing Risks Has Small but Significant Effect on Disease Prediction Models

Our hypothesis expected that competing risk survival models would be enhanced versions of the plain model, and they would thereby generate predictions that differed significantly from the predictions of the plain model. The risks of outcomes that compete, or hinder the observation of the primary disease of interest can lead to models that fit the data improperly. While many risks compete with each of our 18 diseases of interest, the only competing risk we examined was death. Various statistical approaches have been developed to consider competing risks. The approaches we analyzed include the Fine and Gray and multi-state models from the survival package, absolute risk model from the RiskRegression package, and absolute risk model from the iCARE package (Table 1, M1).

The variations of the plain model which considered competing risks were often enhancements to the plain model, as measured by goodness of fit. Specifically, model variations which employed the Fine and Gray or RiskRegression frameworks all generated R^2^ differences that were positive for a majority of diseases. However, the average R^2^ for both these model variations were slightly negative, -6.08E-6 and -1.97E-4 respectively, indicating that their enhancement abilities are modest at best (Table 1, S4.1). The multi-state model variation generated positive R^2^ differences for only 6 diseases, and all four of the iCARE model variations generated positive R^2^ differences for only 1 disease (Table 1, S4.1). Therefore, considerations of competing risk did lead to enhanced models, although the magnitude of this enhancement is slight.

Model variations that considered competing risks generated predictions that were largely equivalent to the plain model. This observed similarity was especially clear for the non-iCARE models, all of which did not require disease incidence estimations (Figure 4A). For example, the average log odds ratio percent difference between the multi-state model and plain model is – 0.13% (SE 0.86%), between the Risk Regression and plain model is 0.69% (SE 0.54%), and between the Fine and Gray and plain model is 0.01% (SE 0.60%) (Figure 4B). However, the value of the polygenic risk score was slightly but consistently elevated in these competing risk models. The average difference in log odds ratio improvement between the multi-state model and plain model was 29.1% (SE 28.1%), between the risk regression and plain model was 12.6% (SE 6.9%) and between the Fine and Gray and plain model was 17.2% (SE 9.7%) (Table S4.5). This pattern of negligible absolute accuracy changes yet evident increases in the predictive contribution of the polygenic risk score was easily seen when examining specific diseases. For example, gout’s difference between multi-state and plain model generated log odds ratio was only 0.3%, but the same model comparison for improved log odds ratio was 442%. Although several diseases, such as rheumatoid arthritis, did buck this trend.

**Figure 4:**
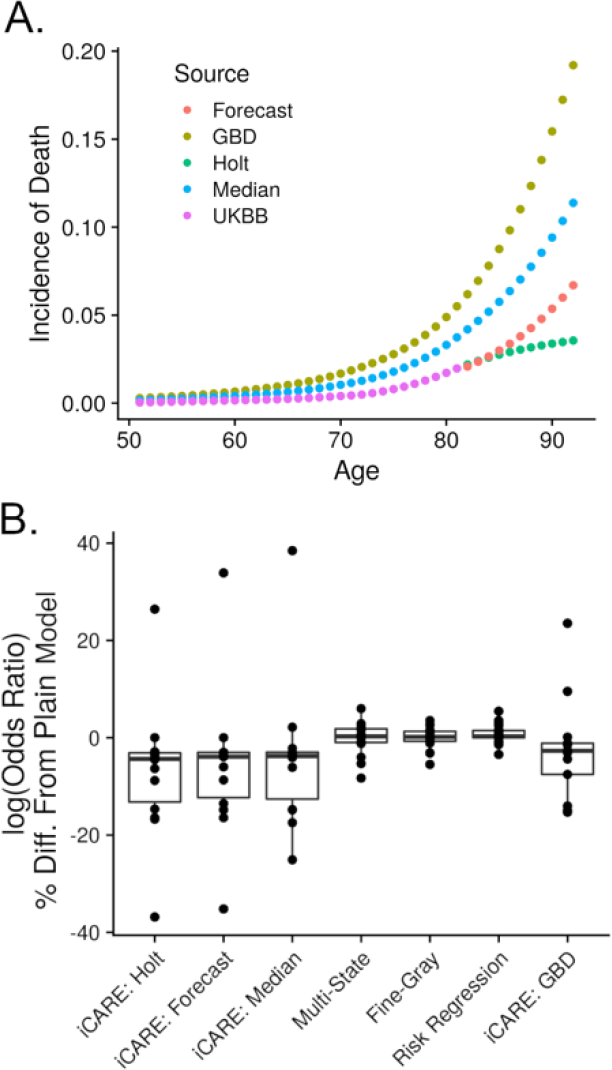
Model Variations that Consider Competing Risks. A: Incidence of death according to various sources and estimates. The GBD incidences are externally estimated from the Global Burden of Disease Study, UKBB are empirically estimated from the cohort under analysis, Median takes the median at each timepoint between GBD and UKBB, Holt extends UKBB by employing Holt’s exponential smoothing, and Forecast are the predictions of a time-series model regressing the UKBB incidence values against the GBD incidence values. B: The percent differences in log odds ratio generated by the model variation indicated and the plain model. Odds ratios are calculated by comparing those in the bottom half of the model’s predictions against those in the top 5%.

We expected competing risks models to have a greater effect on diseases that occur closer to the typical time of death, but found no such association. The correlation between the plain model’s age coefficient and the multi-state model’s absolute log odds ratio difference and the multi-state model’s improvement log odds ratio difference was –0.165 (P = 0.51) and –0.316 (P = 0.20), respectively (Table S4.2). Similar insignificant correlations occurred for Fine and Gray, and Risk Regression analyses. These insignificant results highlight how the trend of a polygenic risk score contributing more to competing risk models as compared to the plain model is relatively modest and undependable.

One clear pattern did emerge within the competing risk models, the iCARE generated predictions were substantially different than the predictions of the plain model. The difference between log odds ratios generated from the iCARE-Holt model and the log odds ratios generated from the plain model was on average –6.8% (SE = 3.2%), with only one disease generating a positive log odds ratio difference (Figure 4B). The difference between the improvement log odds ratios generated from the iCARE-Holt model and the improvement log odds ratios generated from the plain model was 4.23% (SE = 22.0%), with only three diseases generating positive improvement log odds ratio differences. Gout’s difference in log odds ratio improvement between the iCARE- Holt and plain model (295%) highly deviated from expectations and largely swayed the average. A possible explanation for the poor performance of iCARE is its need for estimates of disease and death incidence that exceed the age of a typical UK Biobank participant (Figure 4A). We estimated these disease incidences by combining incidences empirically measured in the UK Biobank and incidences recorded by the Global Burden of Disease Study[13]. Various combination strategies were employed, for example iCARE-Holt represents the extrapolation of the UK Biobank measured incidences with Holt’s exponential smoothing method. However, extrapolation is a notoriously imprecise type of prediction [49] and the Global Burden of Disease incidence values likely do not represent the UK Biobank because of participation bias and likely difference in disease label. For example, the incidence of atrial fibrillation at age 80 is 2.93% according to the UK Biobank and 0.29% according to the Global Burden of Disease Study [13]. Furthermore, the iCARE model is fully parametric, making consistent comparisons to cox proportional hazard models difficult. The strong, unique challenges associated with iCARE models underscore the general difficulty of anticipating the effect when implementing a competing risk model.

### Modified Disease Labels Have Large, Variable Effects on Disease Prediction Models

We hypothesized that modifying the disease labels employed in the plain model would lead to enhanced models whose predictions were substantial different than the predictions of the plain model. The modifications to disease labels were based on the predictions from regularized cox proportional hazard models fit to 2,215 UK Biobank features via the CoxNet algorithm. These features included biomarkers, survey answers, job history, and anthropometric measurements. If the concordance of the regularized model was above 0.75, we compared the model predictions to the plain model’s disease labels. Specifically, we flagged individuals who were either in a top or bottom quantile of the predictions and their disease label differed from what the predictions suggested. These flagged individuals were then either removed from the cohort or reclassified to match the regularized model prediction. The models that utilize these altered disease labels are named according to the quantile of the regularized disease predictions that were inspected, such as 5%, and the action taken, removal or reclassification (Table 1, M1).

Model variations that involved modifications to the disease labels were all enhancements, in terms of goodness of fit, upon the plain model. The R^2^ of the models which reclassified 0.5% of individuals were greater than the R^2^ of the plain models for 15 diseases. The three diseases that did not lead to a positive R^2^ difference include depression, rheumatoid arthritis and type 2 diabetes. No other model variation which modified the diseases labels had fewer diseases which corresponded to positive R^2^ differences (Table 1, S5.1).

When analyzing the predictions of the model variations that modified the disease labels, an initial trend emerged between increased proportions of individuals whose disease label were modified and greater variations in prediction accuracy. Analyses that reclassified individuals in the top or bottom 5% of regularized model predictions generated log odds ratios that differed from the plain model across a range of –73.1% to 122%. When only 0.5% of individuals were considered for reclassification, this range shrank to –31.5% to 60.4% (Figure 5A, 5B). While the reclassification strategies led to models that were just as likely to worsen the plain model’s predictions as improve upon them, the removal strategy appeared to generate consistently better predictions. The average percent difference between the log odds ratio generated by the 5% removal and the plain model was 5.67% (SE 2.38%) (Figure 5A, Table S5.5). However, the 0.5% removal model generated predictions that were largely similar to the plain model, confirming the trend between increased number of modified individuals and greater changes in prediction accuracy.

**Figure 5:**
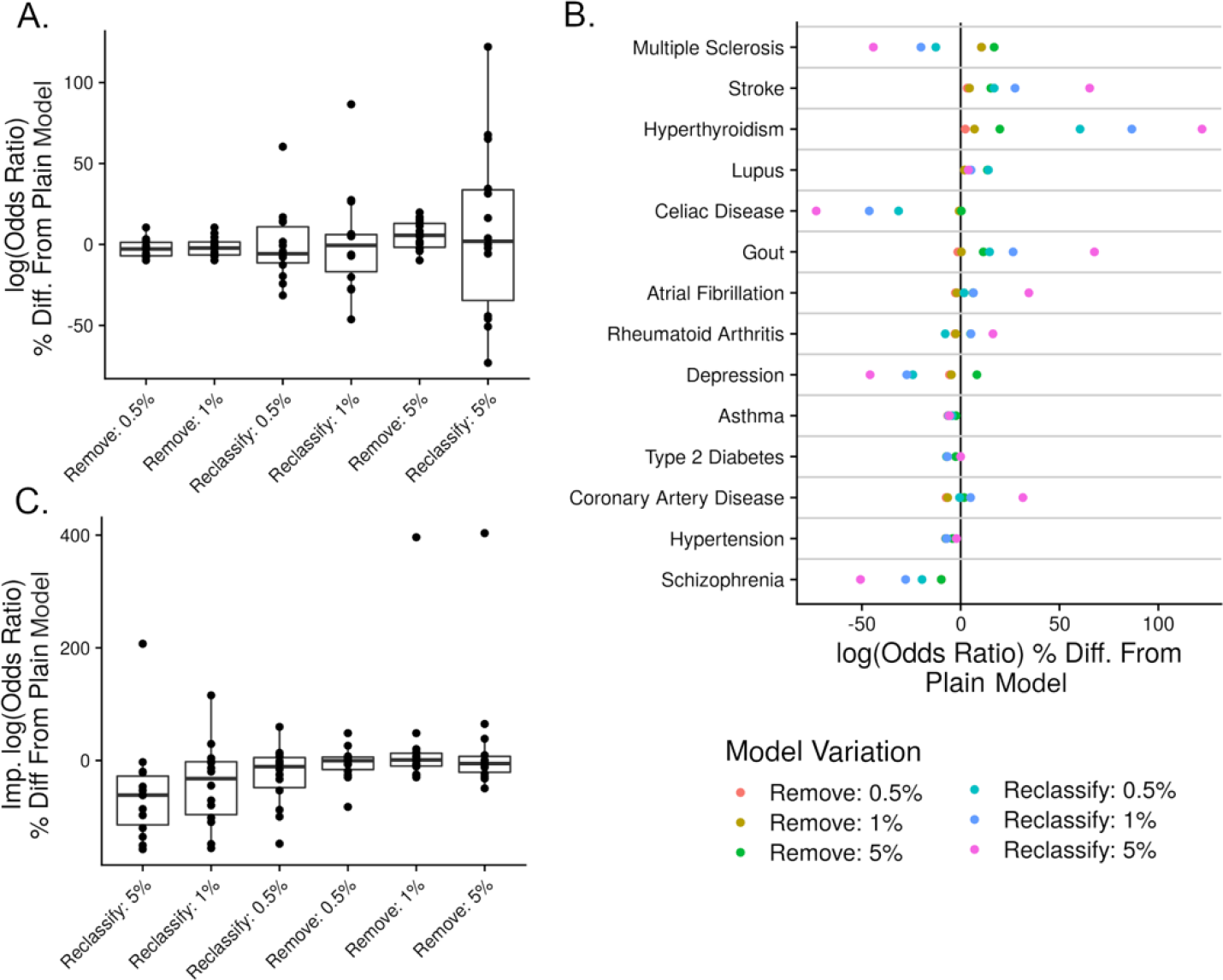
Model Variations that Modify the Disease Labels. A: The percent differences in log odds ratios (calculated by comparing those in the bottom half of the model’s predictions against those in the top 5%) calculated for each disease between a model variation and the plain model, stratified by the model variation employed. B: The values of A stratified by disease. C: Same comparison as in A, except the odds ratio improvement, or the difference in odds ratio computed from the score and base model, replaced the odds ratio directly computed from the score model.

The accuracy improvement brought by the inclusion of a polygenic risk score to a base model was almost uniformly diminished in modified disease label analyses as compared to the plain model. The average percent difference in log odds ratio improvement of the plain model to the 5% reclassification model was –57.5% (SE 30.1%), and to the 5% removal model was –4.1% (SE 8.3%) - when excluding gout. Gout’s difference in log odds ratio improvement between the plain and 5% removal model was 404%, a large value that may stem from both gout’s strong regularized model accuracy (concordance = 0.86) and the large variation in accuracy present across diseases. Diseases that are relatively more difficult to diagnose generated some of the largest decreases in log odds ratio between the 5% reclassification and plain analyses, including lupus [50] (-157%) and celiac disease [51] (-120%) (Figure 5C, Table S5.5). Owing to their diagnostic difficulty, the modified disease labels likely have the greatest chance of bringing these diseases’ labels closer to the ground truth. This interpretation would suggest that contribution of polygenic risk scores to disease prediction models may be overestimated by current, common disease labeling methods.

The trend of modified disease labels decreasing the contribution of a polygenic risk scores to a model was confirmed by concordance metrics, which unlike odds ratios are not sensitive to the small and variable group of individuals designated as high-risk. An initial examination of concordance statistics affirms the variable impact of disease label modifications. The difference in concordances generated by the 5% reclassification model and the plain model varies greatly, from –23.3% to 17.8%, with nearly just as many diseases reporting a positive as a negative concordance difference. However, when examining differences in concordance improvement a more consistent trend emerged. The mean difference in concordance improvement between the 5% reclassification and plain analyses was –67.8% (SE 21.2%). The mean difference in concordance improvement between the 5% removal and plain analyses was greater, although still negative at –8.7% (SE 5.2%) (Table S5.3). The consistency in results across accuracy metrics confirms that disease label modifications cause large variation in prediction accuracy, which often diminish the amount a polygenic risk score effects the predictions of a model.

### Addition of Disease-Specific Covariates to Disease Prediction Models Enhances Their Fit

Our hypothesis expected that variations made to the plain model through the addition of relevant covariates would lead to enhanced models whose predictions differed significantly from the predictions of the plain model. The regularized CoxNet models fit with 2,215 features and utilized in the disease label modification analyses were employed within this collection of model variations. The predictions from the regularized models served as a single, high-impact additional covariate. The coefficients from the regularized model indicated multiple features that could serve as relevant additional covariates (Table 1, M1). The features from each CoxNet model with the ten greatest absolute coefficients were comprised most often by survey answers (32.6%), followed by measurements (21%) and medication usage (21%). Many expected features clearly emerged from the CoxNet models, including insulin use for diabetes, number of days walked ten or more minutes for multiple sclerosis, and bread type consumed for celiac disease (Figure 6A). Although, numerous other unexpected features were included, indicating the difficulty in manually selecting relevant features for a disease prediction model.

**Figure 6:**
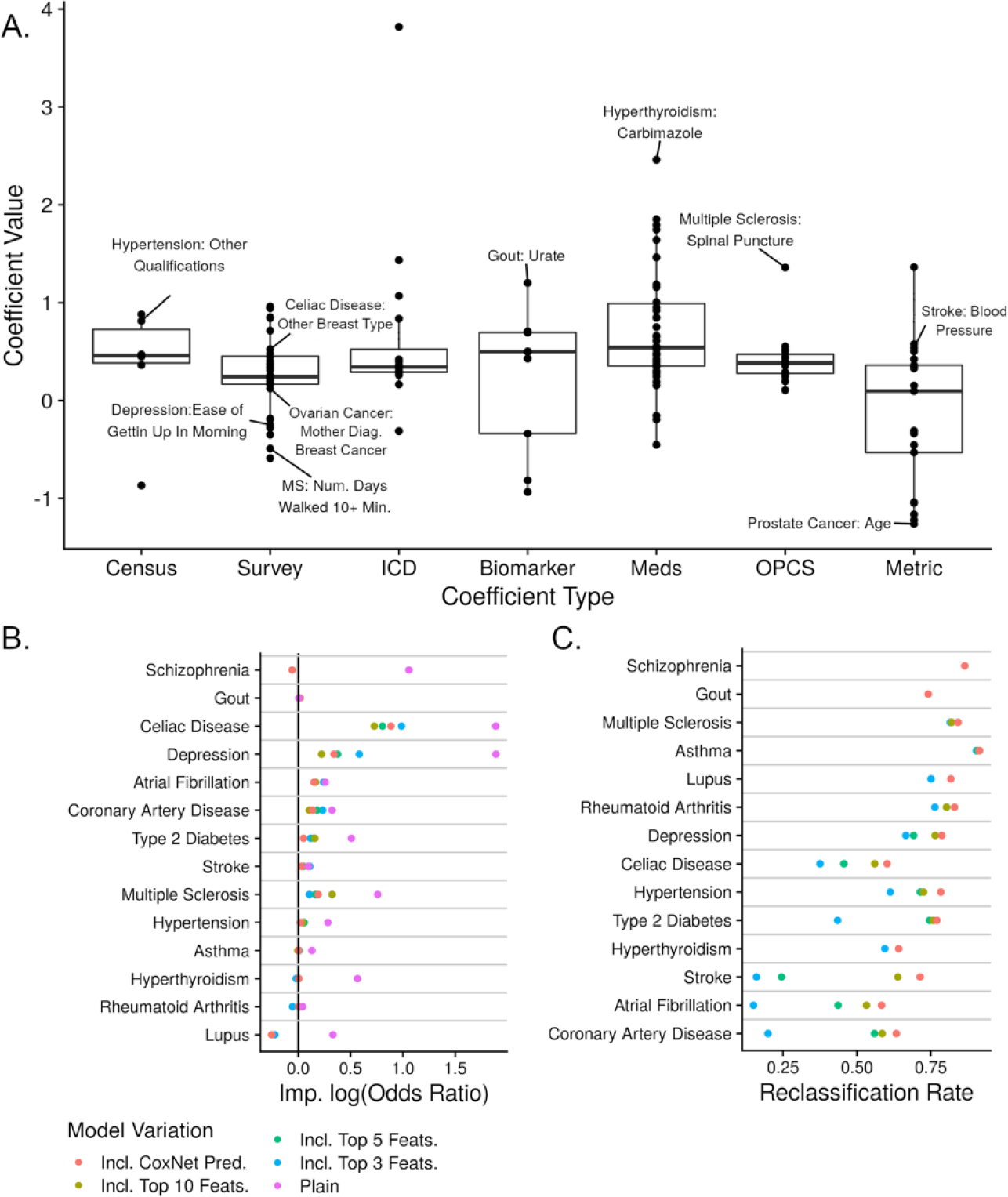
Model Variations that Add Disease Specific Covariates. A: The features with the ten greatest absolute coefficients estimated from each disease’s CoxNet model, organized according to their feature type with certain noteworthy features labeled. B. For each disease, the difference between the model variation indicated and plain model’s improvement log odds ratio. The improvement odds ratio is the difference in odds ratio computed from score and base models, with odds ratio calculated by comparing those in the bottom half of the model’s predictions against those in the top 5%. C: For each disease, the reclassification rates, or fractions of individuals who are no longer considered to be in the top 5% of risk when replacing the plain model prediction with a prediction made from a model variation.

Model variations made through the addition of disease specific covariates were all determined to be enhancements to the plain model in terms of goodness of fit. Even model variations which only included one additional covariate to the plain model generated R^2^ differences that were positive for 13 diseases (Table 1). When the CoxNet model’s prediction was added as a covariate to the plain model, the R^2^ differences were positive for all 18 diseases. The average R^2^ difference of the model variation which included the CoxNet prediction was 0.312 (Table 1, S6.1), which would suggest that on average the CoxNet prediction explains nearly one third of the variance of the average disease under analysis. The model variations which added disease-specific covariates are therefore clearly modelling enhancements.

The accuracies of the additional covariate models were much greater than the accuracies of the plain models. The average difference in log odds ratios corresponding between a model that includes the full CoxNet prediction to a plain model was 128.4% (SE 40.5%). When instead of the CoxNet prediction only the top three features utilized to make the prediction were included, the average difference in log odds ratios only drops to 94.0% (SE 38.6%). Disease specific differences between the log odds ratios generated by the plain and model variation that includes the CoxNet prediction largely varied, with atrial fibrillation having the smallest difference at 18.3% and asthma having the largest difference at 625% (Table S6.5). Although even diseases with small differences in accuracy led to large reclassification rates. For example, with respect to atrial fibrillation 58.4% of individuals considered high risk in the plain model were no longer classified as high risk by the model that included the CoxNet prediction. The average reclassification rate of the model variation that includes the CoxNet prediction was 75.2% (SE 2.8%) (Figure 6C, Table S6.4), indicating how the predictions derived from models with additional covariates largely diverged from the plain model predictions.

The predictive improvement brought by the polygenic risk score was severely diminished within model variations that added disease specific covariates, yet often not completely eliminated. The average difference in improved log odds ratios between the model variation that includes the CoxNet prediction and plain model was –59.2% (SE 28.5%). For five diseases the log odds ratio improvement generated by the model variations that includes the CoxNet prediction was less than 0.01 (Figure 6B, Table S6.5). Similar patterns emerged when examining the statistic of concordance. The average difference in improved concordance between the model variation that includes the CoxNet prediction and plain model was –77.7% (SE 5.9%). For nine diseases the concordance improvement generated by the model variation that includes the CoxNet prediction was less than 0.01 (Table S6.3). While the large decreases in the amount a polygenic risk score effects the predictions of a model may be due to overfitting of the CoxNet model, these results nevertheless indicate that the effects of polygenic risk scores are likely largely overestimated without extensive adjustment.

## Discussion

The results appear to confirm our hypothesis: many of the variations made to the plain model fit the data better and generated predictions which differed significantly from the plain model’s predictions. The plain model specifically is a cox proportional hazard model that regresses age, sex, top ten genetic principal components and a polygenic risk score against time in the study until an electronic heath record derived disease label. The variations made to this plain model fall under five facets of modelling: representation of age, censorship, competing risks, disease labels and additional, disease-specific covariates. Model variations made in response to all five of these areas created massive changes in risk classifications. For example, at least one of the model variations generated a reclassification rate greater than 33% for each disease, implying that if three individuals are identified as high risk under the plain model at least one, on average, will in fact be low risk if an alternative model is utilized. These differing predictions had widely variable effects on both the accuracy of the overall model and the increase in accuracy that occurs when a polygenic risk score is added to a model. In some instances, the model variations, which were intended to be an improvement upon the plain model, fit the data worse. On average, 13 of the 26 model variations made to the plain model were found to decrease the goodness of fit, as measured by Royston’s R^2^. However, we still consider these eight model variations to be potentially useful as literature sources claim they reduce biases or assumptions held in the traditional, plain model. To determine the veracity of these claims we will now discuss the potential benefit and possible caveats underlying each of the five areas of model variations.

First, model variations were made regarding the representation of age. In the plain model, age is represented by a single covariate of age at time of analysis. This approach assumes a linear relationship between age and disease. However, previous investigations have shown this is not always true. Age often has complex, non-linear association to disease risk, as evidenced by the inclusion of square and interaction terms in many genome-wide association analyses [17]. Other than the inclusion of complex model covariates, a common way to handle these biases is shifting the cohort composition so that the effect of age upon disease can be relatively removed [18].

The results from such age-matched cohorts indicate that the contribution of polygenic risk scores to models are underestimated in plain analyses. Although, matched-cohort studies also carry a host of biases [52]. Therefore, while age can certainly be better modeled in polygenic risk score analyses, and doing so may increase the effect of the scores, further investigations are needed to ensure all meaningful biases have been removed.

Second, the right-censorship methodology of the plain model was modified. A right censorship framework tracks each individual from the time they entered the study until they either experienced the disease or were censored. The downside of such a right censored approach is that individuals who enter the study at relatively later times appear less likely to develop disease owing to their decreased exposure time. This participation bias is shared by several other high profile examples, such as a 1968 study of radiation exposure on Hiroshima atomic bomb victims [53], and has been clearly identified in epidemiological-focused publications [20, 54]. While delayed-entry reduces this bias, it raises the question of time-scale. Several investigations reach conflicting conclusions on whether study-time or age-time is the preferred time-scale [22,55,56]. Our results indicate that the age-time scale allows for more flexible, non-parametric modeling of age which increases the contributions of polygenic risk scores to their respective models. Further model variations may incorporate left-censorship, which is able to recognize individuals who are diagnosed at an ambiguous time before they were assessed. Such model variations may lead to wholly new trends between censoring methods and polygenic risk score effects.

Third, competing risks were incorporated into the plain model. In the plain model every individual is assumed to eventually develop the disease as right censorship only implies that the time of diagnosis is greater final time point of the study [23]. However, once an individual dies their chance of developing any disease in the future becomes completely irrelevant. To represent this fact in survival models the competing risk of death can be included, which has the added benefit of generating clinically relevant, absolute risk predictions [25]. Although, participation bias in the UK Biobank [57] explicitly limits the time-extent of such predictions. Possible corrections to these biases, along with the use of improved multi-state schemes [58], may further exacerbate the accuracy differences between competing risk and plain models.

Fourth, the disease labels employed within the plain models were modified. We suspected that survey and electronic health record derived disease labels may be imperfect because some individuals with a disease diagnosis also held a highly contradictory feature. For example, 148 individuals with diabetes report HbA1c levels less than 35 mmol/mol, even though 48 mmol/mol is traditionally the diagnosis cut-off [59]. Several recent investigations have proven the practicality of using a wide range of diverse features to form accurate disease predictions that are robust to any single error in the electronic health records [28, 29]. However, these predictions are still far from perfect. Any results drawn from disease labels modified according to these predictions are therefore inherently unreliable. The nebulous conclusion drawn from our results suggests polygenic risk scores are highly sensitive to the disease labels they are compared to, leading to easy overfitting which overestimates their predictive ability. Improved alternative disease predictions may alter this understanding.

Fifth, relevant, disease-specific covariates were added to the plain model. Without all salient covariates in a model the polygenic risk score may act as a proxy for other features, thereby developing an inflated effect size. Manually selected sets of relevant covariates could easily miss important risk factors or conversely may include features that do not have a true, useful association to the disease of interest. We therefore implemented an unbiased feature selection approach. As expected, inclusion of additional covariates in the disease prediction model increased overall accuracy and decreased the value of the polygenic risk score. The polygenic risk score still contributed to nearly all disease prediction models. Although, our alternative predictions may be overfit or underperforming, leading to an underestimation or overestimation of each score’s effect, respectively. Further investigations are needed to determine proper, unbiased model adjustments.

Along with the limitations that are specific to the five areas of modelling assumptions, other limitations exist that apply to the entirety of the investigation. First, while we examined a relatively wide variety of model variations, there are many other possible variations that could plausibly mitigate the assumptions and biases of the plain model. The fit and accuracy of these unexamined variations is unknown. Second, we only analyzed individuals of “White British ancestry”, limiting the direct impact of our results to like individuals [3], although we expect the general epidemiologic principles reported to be ancestry insensitive. Third, several of the polygenic risk scores were derived from small and simple sets of genetic variants. Advances in constructing variant sets would likely improve the amount a polygenic risk score effects a model’s predictions and could possibly alter the trends drawn from our results. Various other assumptions regarding participation bias and data authenticity have also been made [57,60,61].

Despite these limitations, the predictions of the model variations consistently differ from the plain model’s predictions. In certain areas, the differences align to form noteworthy and explainable patterns that ought to be mediated in future analyses. The contribution of the polygenic risk score to its respective model tends to be increased by variations to the representation of age and censorship and tends to be decreased by variations to disease labels and the addition of disease-specific covariates. With terrific amounts of variance contained in these trends any future investigation cannot take for granted that their polygenic risk score will be similarly affect by these model variations. These future investigations must therefore employ the model variations and assess their effects upon their specific set of circumstances. We would expect that the model variations would have beneficial effects upon the predictions, as literature supports the ability of the variations to reduce problematic assumptions and biases in the models. This expectation has been confirmed regarding the model variations which fit the data better than the plain model. Without implementing these enhanced model variations, polygenic risk scores may currently be improperly discarded or exulted in preliminary research and one day may even provide a distinctly inferior risk prediction to a patient facing a difficult clinical choice. With little to no downside, we believe that future polygenic risk score analyses should evaluate models with all possible variations.

## Methods

### Materials

The UK Biobank, a large, prospective, general health-study, provided all necessary genotypic and phenotypic data[62]. A total of 502,419 individuals were initially assessed by the UK Biobank from 2006 to 2009, from which 408,045 individuals of White-British ancestry who met standard quality-control procedures were retained for analysis[63]. From these individuals, genotypic information comprised 6,939,064 imputed genotypic variants which met standard quality control metrics[8]. Phenotypic information was defined by both self-reports of disease and ICD/OPCS hospital codes (Table M2, M3). Individuals whose disease label occurred before their date of assessment within the UK Biobank was removed from the analysis of that specific disease. Environmental, clinical and demographic information was drawn from survey answers, blood test results, and anthropometric measurements. Supporting information used to further describe each UK Biobank individual included the 2011 United Kingdom local super output area census data, ascribed to each individual based on their reported home location, and expected age-specific disease incidence as recorded within the Global Burden of Disease investigation[13].

The UK Biobank was accessed through application #47137. The UK Biobank has approval from North West Multi-Centre Research Ethics Committee. All participants gave written, informed consent.

### Polygenic Risk Scores

Polygenic risk scores were calculated by combining genetic variant weights extracted from the PGS Catalog and the genotypes of all individuals in the UK Biobank (Table M4). The sets of genetics variant weights were quality-controlled such that the constituent variants were non- ambiguous, within the UK Biobank imputed genotypes, and properly allele-flipped. Next, polygenic risk scores were computed by PLINK[64]. We selected the 18 variants sets as they spanned unique diseases, and each contained over 50 variants.

### Non-Genetic Disease Predictions

Disease risk models that utilized a large, diverse dataset were utilized with model variations that modified disease labels and model variations that added disease-specific covariates. The dataset for these predictions were carefully quality controlled and imputed (Table M5, M6). The models were fit with the CoxNetSurvivalAnalysis function from the sksurv package, which is a regularized model that is penalized for the number of non-zero coefficients it contains. Greater information concerning the construction of the dataset and the application of the CoxNet function can be found in the supplemental methods.

### Statistical Analysis

The polygenic risk scores for all 18 diseases were chiefly analyzed by comparing a series of paired cox proportional hazard models. The base model in the pair contained the covariates of age, sex, and top ten genetic principal components, whereas the score model contained all the covariates of the base model and the polygenic risk score. A plain model type regressed the aforementioned covariates against time from initial assessment to disease, while the 26 variations of the plain model, or model variations, changed some aspect of the plain model with the intent of reducing assumptions or biases held within the plain model. Each model could either be plain or a variation, and score or base – terminology that is employed throughout the investigation. Comparisons between score and base model statistics were termed statistical improvement. Comparisons between plain and variation model statistics were termed differences.

Variations to the plain model were made in five areas: representation of age, censorship, competing risks, disease labels and additional covariates. Each area included multiple model variations, for example the competing risk area generated Fine and Gray models, multi-state models, and absolute risk models fit with the RiskRegression package. A short description of each model variation is provided in table 1. Details on how each model was computationally created is provided in table M1. Additional descriptions and motivations explaining how each model variation reduces the assumptions of the plain model are provided in the introduction, relevant results section and discussion.

Models were fit upon a training phase of data and assessed on the testing phase through computation of goodness of fit metrics, model coefficients, concordance, reclassification rates, and odds ratios. Specifically, reclassification rates were defined as the fraction of individuals who were no longer in a high-risk quantile specified by an individual’s final time-point cumulative hazard when shifting from the simple model to another. Odds ratios were computed by forming a contingency table in which low and high-risk individuals held final time-point cumulative hazards below the 50^th^ percentile and above the 95^th^ percentile, respectively. A model variation was determined to be an enhancement upon the plain model if it fit the data better for more than half of the diseases assessed. Goodness of fit was measured by Royston’s R^2^.

The R coding language version 3.6 (with packages survival, icare, tidyverse, epitools) and the python coding language version 3.6 (with packages sklearn, sksurv, pandas and numpy) carried out all computations. All scripts are readily available at https://github.com/kulmsc/prs_surv_models, all acronyms employed are described within Table M7, and additional methodological descriptions are available in the supplementary.

